# Leaf gene expression trajectories during the growing season are consistent between sites and years in American beech

**DOI:** 10.1101/2022.12.17.518988

**Authors:** U. Uzay Sezen, Jessica E. Shue, Samantha J. Worthy, Stuart J. Davies, Sean M. McMahon, Nathan G. Swenson

## Abstract

Transcriptomics provides a versatile tool for ecological monitoring. Here, through genome-guided profiling of transcripts mapping to 33,042 gene models, expression differences can be discerned among multi-year and seasonal leaf samples collected from American beech trees at two latitudinally separated sites. Despite a bottleneck due to post-Columbian deforestation, the SNP-based population genetic background analysis has yielded sufficient variation to account for differences between populations and among individuals. Our expression analyses during spring-summer and summer-fall transitions for two consecutive years involved 4197 differentially expressed protein coding genes. Using *Populus* orthologs we reconstructed a protein-protein interactome representing leaf physiological states of trees during the seasonal transitions. Gene set enrichment analysis revealed GO terms that highlight molecular functions and biological processes possibly influenced by abiotic forcings such as recovery from drought and response to excess precipitation. Further, based on 324 co-regulated transcripts, we focused on a subset of GO terms that could be putatively attributed to late spring phenological shifts. Our conservative results indicate that extended transcriptome-based monitoring of forests can capture diverse ranges of responses including air quality, chronic disease, as well as herbivore outbreaks that require activation and/or downregulation of genes collectively tuning reaction norms maintaining the survival of long living trees such as the American beech (*Fagus grandifolia*).

## Introduction

Anthropogenic climate change is affecting the phenology of trees, with important implications for the future of forest health [1, 2]. Trees over their long lifespan experience a multitude of severe conditions, but climate change is shifting essential phenological events and intensifying many of the environmental deviations. These deviations can exceed the boundaries of reaction norms trees have evolved [3–8]. For temperate trees, changes in the arrival of spring, and shifting precipitation regimes can affect the phenological scheduling of vital biological events such as vessel activation [9], the acquisition of soil nutrients [10], flowering [3, 11–13], leaf flushing [14–18], xylogenesis [19], seeding and fruiting [20–22], and senescence induced nutrient re-uptake from leaves [16, 23–26]. Ultimately, the resulting evolutionary dynamics will redefine competition and reshape species ranges [18, 27, 28]. Functional responses to these climatic changes, however, are complex and difficult to demonstrate against the background noise of ecological variation. Because genes fundamentally drive all these mechanisms, gene expression might offer a tool towards identifying patterns to advance our understanding of the mechanisms of organismal response to climate change through phenology [29–33].

Leaves are an important component of individual-level performance forming an interface between the phenotype and the environment. Phenotypic variation is an interplay of the genotype and gene expression. Yet, little is known about the relative significance of these two constituents of the phenotype. Fine-scale transcriptomic assays of leaf function of individual trees through space and time are scarce, rarely cover an entire growing season, and/or have not been carried out at more than one site [34–36]. There is an urgent need for determining the drivers of tree leaf function especially among trees growing in unmanaged natural forest stands. The present study aims to fill this gap through a comprehensive transcriptomic assay of leaf gene expression through space and time.

Transcriptomic analysis captures expressed genes in a tissue at the time of collection and can provide detailed sequence information allowing the quantification, comparison, and accurate interpretation of phenotypes along a phenological sequence. Leaves, in particular, are organs that have evolved adaptations to sense the environment and acclimate in order to maintain homeostasis. Leaves help regulate whole plant performance by balancing the uptake of carbon for photosynthesis and hydraulic flow. Measuring gene expression in leaves, therefore, offers a way to probe into developmental and plastic responses throughout the growing season, linking the influences of climate shifts to the phenotypic response of the trees. Importantly though, no two growing seasons are alike making quantification of gene expression in leaves across space and time vital to further insight into plant function and status under different environmental conditions and phenological stages [3, 37–39].

The timing of seasonal transitions has been shown to differ among years and sites, driven by changes to temperature and precipitation patterns. Unfortunately, these and other climatic variables are expected to be further altered by climate change and become less predictable potentially leading to mismatches and disruptions in pollination services and trophic interactions such as herbivory, predation and parasitizitation [3, 11–13, 40, 41].

American beech (*Fagus grandifolia* Ehrh.) is a common tree, the sole member of the genus naturally found in the northeast United States and a characteristic feature of older growth forests. *Fagus* species are known for their responsiveness to the environment, with an ability to grow slowly when resources are not available, and quickly when conditions reverse such as increased light after gaps form in the canopy or soil humidity is restored [42]. The genus is particularly sensitive to drought as compared to other co-occurring hardwood species [43]. In this study, we investigated patterns of leaf gene expression in two populations of *F. grandifolia*: one in Maryland, USA and the other in central Massachusetts, USA. After collecting two years of expression data, three times each year, we quantified how gene expression differs across sites and seasons. We leveraged information regarding the up- and down-regulation of genes in each leaf sample collected from the same trees, we asked the following questions: 1) Does leaf gene expression vary with respect to year, site, and season? 2) Can site or annual differences in expression be explained by abiotic factors? 3) Do the identities of the differentially expressed genes suggest mechanisms that control phenology?

## Results

### Genome alignments and background genetic analysis

Alignment rates of trimmed reads to the draft American beech genome were high. Close to 95 percent of the samples had reads aligned with a rate more than 80% (SI Fig. S3, Table S1). Alignments spanned 33,042 gene models corresponding to 27,226 genes in the draft *F. grandifolia* genome (unpublished draft American beech genome sequence). Although the forests of the US east coast are a product of large-scale second-growth regeneration after large-scale deforestation, populations of beech trees at the two ForestGEO FDP sites namely, Smithsonian Environmental Research Center (SERC) and Harvard Forest (HARV) displayed enough genetic variation to distinguish them based on SNP variants (Fig. S4). A pairwise comparison of SNP profiles of 40 trees generated a kinship pattern where the within population relatedness exceeded that of between population (Fig. S4). Further, a Kendall’s rank correlation of association on kinship coefficients had positive values for within population comparisons reflecting local founder effect (Fig. S4, Table S1). In both populations, there are few distantly related individuals that stand out with negative correlations (Fig. S4). On the other hand, correlations were negative using between population comparisons reflecting genetic effect of latitudinal separation by more than 640 km (Fig. S4, Table S1).

### Does leaf gene expression vary more with respect to year, site or season?

The first two axes of a Principal Component Analysis (PCA) on the leaf gene expression data showed that the three seasons were transcriptomically distinct (Fig. 1). Site differences were not very pronounced and mostly followed a parallel expression trajectory, except in the spring 2018, which shows the most separation between the research sites (Fig. 1). The yearly dynamics of each season showed a recognizably linear PC1 distribution consistent across both populations. The first two components together represented 73.5 % of the total variation. The spring samples were separated along both PC axes to a greater extent and represented the largest magnitude of change during seasonal transitions in both years (Fig. 2). Summer and fall samples together formed tighter clusters, but yearly dynamics were still distinguishable (Fig. 1). Once summer was reached, the expression pattern of genes remained relatively unchanged into the fall. Specifically, summer and fall samples largely overlapped especially in 2017 (Fig. 1). Fall 2017 samples from SERC overlapped with their summer counterparts representing a sustained gene expression pattern along summer-fall transition (Fig. 1 unshaded orange and blue circles). Overall, the 2018 growing season casted a wider gene expression space while 2017 represented a more contracted parallel transcriptional activity in both sites (Fig. 1).

**Figure 1.**
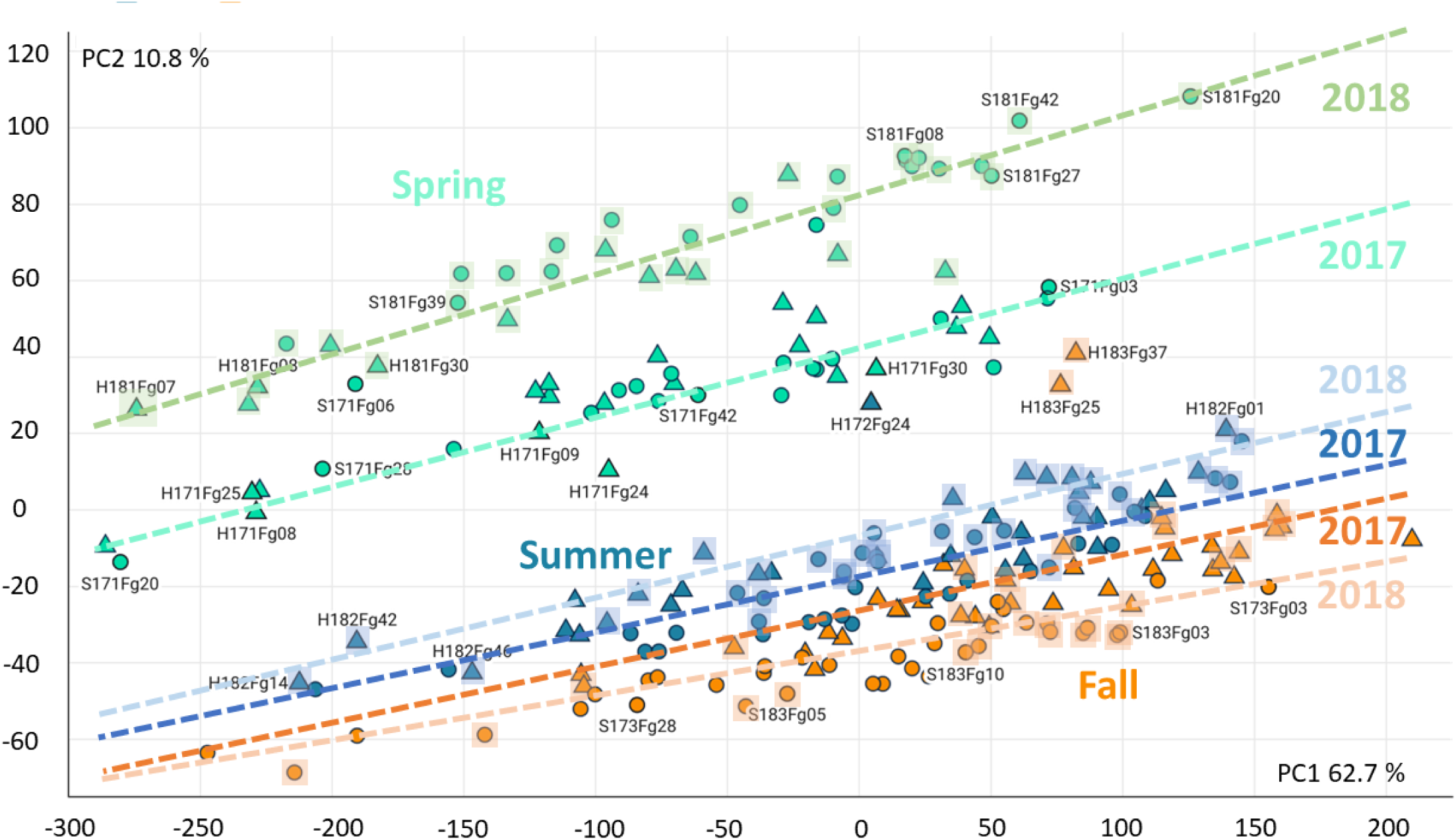
A PCA of differentially expressed genes in 222 *F. grandifolia* samples collected from 40 trees during Spring, Summer and Fall of 2017 and 2018 from HARV (triangles) and SERC (circles). The PCA used singular value decomposition with imputation and includes the top 11.500 most variable loci that were ln(x) transformed and Pareto scaled. Shaded regions represent 2018 samples. The PC1 and PC2 represent 73.5 % of the total variation. Dashed lines were inserted manually to show the approximate trends across years.

**Figure 2.**
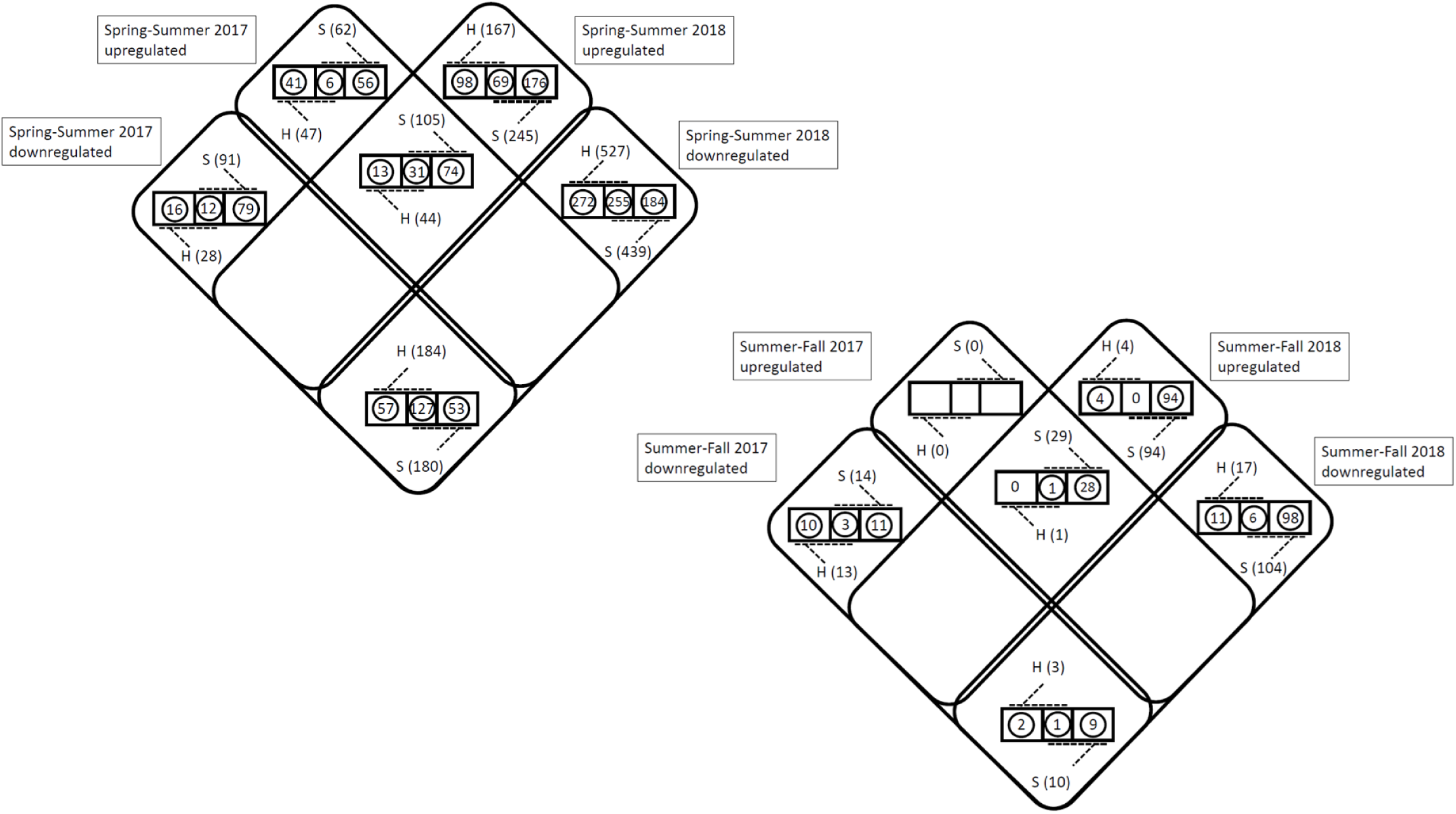
A breakdown of differential and co-expression patterns during spring-summer, summer-fall transitions at two sites along two consecutive years. Sites are coded as H (HARV) and S (SERC).

### Mapping differential gene expression on an interactome

The total number of differentially expressed transcripts corresponded to 4197 American beech protein models (SI Dataset S1). A search for these proteins against the black cottonwood (*Populus trichocarpa*), NCBI taxon identifier 3694) matched 2813 orthologs. Of the *Populus* orthologs, a subset of 2421 had a hit in the STRING database generating an interactome network with 2185 nodes and 3231 edges. The largest patch consisted of 1280 nodes and 3185 edges (Fig. S10). The network contained 1329 singleton nodes (Fig. S10, singleton nodes not shown for brevity). Consistent with the PCA results, overall the highest magnitude of change took place during the spring-summer transitions during both 2017 and 2018 (Fig. S10).

### Gene expression during spring-summer transitions

The trees at the two sites consistently co-upregulated 31 and co-downregulated 127 transcripts (*Populus* orthologs) in both years during spring-summer transitions (Fig. 2, Venn diagram left). In HARV, 13 transcripts were upregulated exclusively in both years (Fig. 2, Fig. S10 green fill with red border). The number of transcripts that were upregulated solely in SERC for two consecutive years was 74 (Fig. 2, Fig. S10 yellow fill with red border). In 2017, SERC had 56 and HARV had 41 transcripts upregulated in a site-specific manner. In the following year, the site-specific expression levels were roughly two-fold higher in HARV with 98 transcripts and three-fold in SERC with 176 transcripts (Fig. 2). In parallel, site-specific downregulation in 2018 was 16-fold higher in HARV with 272 transcripts (Fig. 2). SERC had two-fold higher downregulated site-specific expression with 184 transcripts (Fig. 2). Therefore, spring-summer transition in 2018 demonstrated the largest differential gene expression activity in parallel at both sites. There were 127 transcripts co-downregulated in both sites and years (Fig. 2). HARV had 57 transcripts repeatedly downregulated during spring-summer transitions in both years (Fig. 2). SERC had 53 transcripts consistently downregulated in two successive years (Fig. 2).

### Gene expression during summer-fall transitions

Gene expression patterns during summer-fall transitions at the two sites did not differ as much as observed in Spring-Summer and therefore it was possible to represent differential expression from both sites in a single panel at a given year (Fig. 2, Venn diagram right). There was a single *Populus* ortholog each for repeatedly upregulated (Acid Phosphatase 1) and downregulated (Nuclear Fusion Defective 4) transcripts found in both years at two sites (Fig. 2). Strikingly, there were no upregulated transcripts in either of the two sites in summer-fall 2017. In 2017, the two sites experienced a relatively homogeneous summer-fall transition with a moderate amount of 10 and 11 downregulated transcripts with only three co-downregulated transcripts (Fig. 2). SERC appears to have a more dynamic summer-fall transition beginning with 28 *Populus* orthologs upregulated as opposed to none in HARV (Fig. 2). There were 94 *Populus* orthologs specifically upregulated in SERC while in HARV there were only four (Fig. 2). No co-expressed orthologs were shared between the two sites in 2018 (Fig. 2). Downregulated transcripts particular to SERC in 2018 corresponded to 98 *Populus* orthologs almost 10-fold higher than that of HARV with 11 (Fig. 2). Co-downregulated *Populus* orthologs in both sites amounted to six (Fig. 2).

### Enriched GO terms during spring-summer transitions

Up and downregulated transcripts reflected both latitudinally parallel and divergent patterns in enriched Gene Ontology (GO) terms including biological process (BP), molecular function (MF), cellular compartment (CC) and KEGG (Kyoto Encyclopedia of Genes and Genomes) suggesting a strong environmental influence on response of both forest sites (Fig. S8). In 2017, upregulated transcripts in HARV were grouped under 8 terms belonging to sugar and carbohydrate transport activity (Fig. S8). During the wet growth season of 2018, trees in both sites showed a parallel downregulation of transcripts encompassing 23 cell wall and membrane biosynthetic terms including very long chain fatty acid biosynthesis that control suberin and epicuticular wax quality as a response to abiotic stress (Fig. S8). A list of top 20 differentially expressed transcripts with their draft genome accessions and annotations are provided for all seasonal transitions (Table S1).

### Genes possibly involved in phenology of late-stage leaf expansion affected by delayed spring

A total of 324 transcripts were simultaneously co-regulated (69 upregulated, 255 downregulated) at both sites as a potential response to delayed spring leaf expansion during the spring to summer transition in 2018 (Fig. 2). These transcripts may potentially be controlling the phenology of delayed leaf expansion enriched with 6 GO terms pertaining to biological processes, 5 terms of molecular function and 11 terms involving cellular components. Additionally, these transcripts were enriched for 5 KEGG metabolic pathways (Table 1). Enriched terms related to the membrane (GO:0031224, GO:0016021, GO:0005886, GO:0016020) and metabolic pathway of fatty acid elongation (pop00062) are associated with phenological transitions (Table 1). A full list of STRING identities for these transcripts and their annotations are included in the supplementary file (Table S1).

**Table 1.** STRING output of enriched GO terms from 324 co-regulated transcripts possibly involved in delayed spring leaf expansion phenology in *F. grandifolia*. Permalink to the analysis: https://version-11-5.string-db.org/cgi/network?networkId=bWUhWsqb5LZt

| Biological Process |  |  |  |  |
| --- | --- | --- | --- | --- |
| GO-term | description | count in network | strength | false discovery rate |
| GO:0098754 | Detoxification | 10/238 | 0.73 | 0.037 |
| GO:0071669 | Plant-type cell wall organization or biogenesis | 11/297 | 0.67 | 0.037 |

**Table 1 continued from previous page**
|  |  |  |  |  |
| --- | --- | --- | --- | --- |
| GO:0071555 | Cell wall organization | 24/716 | 0.63 | 1.91E-05 |
| GO:0071554 | Cell wall organization or biogenesis | 28/894 | 0.6 | 8.34E-06 |
| GO:0005976 | Polysaccharide metabolic process | 17/658 | 0.52 | 0.037 |
| GO:0005975 | Carbohydrate metabolic process | 30/1581 | 0.38 | 0.0189 |

**Molecular Function**
| GO-term | description | count in network | strength | false discovery rate |
| --- | --- | --- | --- | --- |
| GO:0004601 | Peroxidase activity | 8/161 | 0.8 | 0.0313 |
| GO:0016747 | Transferase activity | 16/581 | 0.55 | 0.0285 |
| GO:0046906 | Tetrapyrrole binding | 17/646 | 0.53 | 0.0285 |
| GO:0020037 | Heme binding | 16/607 | 0.53 | 0.0285 |
| GO:0003824 | Catalytic activity | 159/14494 | 0.15 | 0.00044 |

**Cellular Component**
| GO-term | description | count in network | strength | false discovery rate |
| --- | --- | --- | --- | --- |
| GO:0009505 | Plant-type cell wall | 9/195 | 0.77 | 0.004 |
| GO:0048046 | Apoplast | 15/508 | 0.58 | 0.0025 |
| GO:0005576 | Extracellular region | 60/2170 | 0.55 | 3.70E-14 |
| GO:0005618 | Cell wall | 19/766 | 0.5 | 0.0025 |
| GO:0005794 | Golgi apparatus | 26/1479 | 0.35 | 0.0149 |
| GO:0031224 | Intrinsic component of membrane | 96/7034 | 0.24 | 9.49E-06 |
| GO:0016021 | Integral component of membrane | 90/6767 | 0.23 | 4.27E-05 |
| GO:0071944 | Cell periphery | 80/6039 | 0.23 | 0.00026 |
| GO:0005886 | Plasma membrane | 65/5314 | 0.19 | 0.0179 |
| GO:0016020 | Membrane | 130/11031 | 0.18 | 3.22E-05 |
| GO:0110165 | Cellular anatomical entity | 272/29493 | 0.07 | 3.14E-05 |

**KEGG pathway**
| pathway | description | count in network | strength | false discovery rate |
| --- | --- | --- | --- | --- |
| pop00062 | Fatty acid elongation | 5/40 | 1.2 | 0.0017 |
| pop00520 | Amino sugar and nucleotide sugar metabolism | 7/154 | 0.76 | 0.0094 |
| pop00940 | Phenylpropanoid biosynthesis | 8/206 | 0.7 | 0.0094 |
| pop01110 | Biosynthesis of secondary metabolites | 26/1446 | 0.36 | 0.0044 |
| pop01100 | Metabolic pathways | 42/2605 | 0.31 | 0.0013 |

## Discussion

There have been a few transcriptomic time course studies produced to understand multi-year, multi-season dynamics in plants [19, 34, 35, 44, 45]). These works have largely focused on a single site and/or a single growing season with a single study on oak representing a rare exception spanning a decade in a multiple site setting [33]. Here, we have produced a detailed study of leaf gene expression in *Fagus grandifolia* in two distinct populations in the eastern United States through the growing season in multiple years. Despite the climatic differences in years and sites, that we describe below, and the potential for a high degree of variation in leaf gene expression profiles, it is notable that several emergent and consistent patterns were found. First, gene expression profiles significantly change between the spring and summer components of the growing season. Second, unlike the spring-summer transition, gene expression profiles are generally consistent from the summer leading into the fall. Third, and perhaps most surprisingly, these results were consistent across years and across genetically distinct populations separated by hundreds of miles and several degrees of latitude. These results indicate that, while leaf gene expression profiles are dynamic, they are largely predictable across seasons, sites and years. In the following, we discuss the results in detail beginning with information regarding the sampling conditions across seasons and years.

Gene expression is, typically, considered to be highly labile and the quantification of gene expression in the wild may be expected to lead to incredibly noisy signals due to the variable ecological contexts experienced by a tree in space and time. For example, the eastern United States experienced a substantial drought period beginning in 2016, the year prior to our sampling, lasting into the spring of 2017 (SI Fig. S5). Thus, during the 2017 growing season, trees at the HARV site were simultaneously recovering from drought while also dealing with the ongoing impacts of beech bark disease (SI Fig. S5, SI Fig. S7). In 2018, spring arrived late compared to the previous year (Fig. 3), and many states across the east coast had record breaking rainfall (SI Fig. S6, SI Fig. S5, SI Fig. S11-14). Specifically, May 2018 to May 2019, the continental United States experienced the most abundant rainfall recorded in 124 years (SI Fig. S6, SI Fig. S13-14).

**Figure 3.**
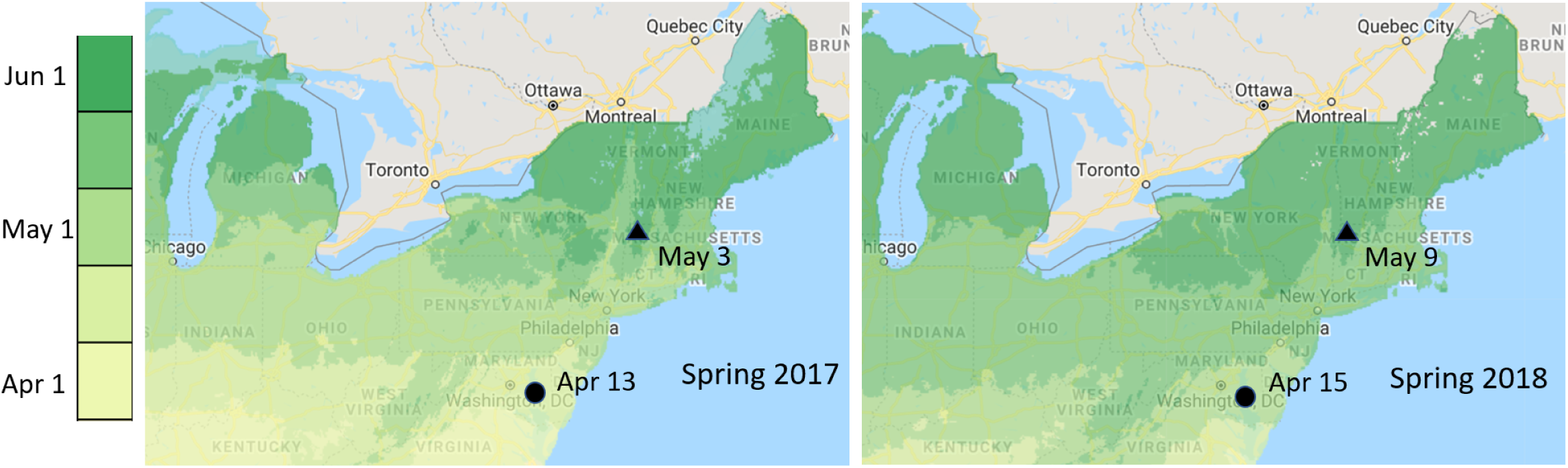
Phenology of Spring arrival comparison based on first bloom between 2017 and 2018. In 2018, the spring arrived late along the entire Northeastern US. The visualization is an annual representation of the days of year that the requirements for the first bloom Spring Index were met, averaged for Red Rothomagensis lilac, Arnold Red honeysuckle and Zabelii honeysuckle; available since 1981, calculated using PRISM Tmin and Tmax data at a particular location. First leaf out dates for the genus Fagus have been indicated next to the sites. Data from US National Phenology Network (USA-NPN) and NEON.

Taking these yearly and site variations into consideration, one would expect that the beech trees in this study would have highly variable gene expression profiles within and between sites and years. As our results show, the combined effect of abiotic factors such as late spring and precipitation can generate a discernable molecular phenotype between the same seasons of consecutive years. The pattern is visible among Spring 2017 - Spring 2018 samples of both sites separated along the PC2 (Fig. 1). Nearly each spring has a distinct signature with a latitudinally parallel trajectory along both sites (Fig. 1). That is, in each season within a given year, our trees had a transcriptomic pattern following a year-specific trajectory (Fig. 1).

Spring samples casted a wider ordination footprint compared to summer and fall (Fig. 1). This means the largest differential gene expression change is observed during the spring-summer transitions. The vigor observed during the spring-summer transition is evident in the mapping of the differentially expressed transcripts on the *Populus* interactome through their orthologs (Fig. S10). The arrival of spring is a continually moving target and increasingly becoming so with climate change [1, 2, 11–13, 46]. Therefore, understanding spring-summer growth dynamics is important since global warming is expected to induce phenological shifts in flowering and leaf-out timings of most trees and affect mortality [3, 4, 7]. In parallel, we particularly paid attention to a set of 324 co-regulated transcripts in both study sites that could possibly be involved in controlling the phenology of delayed leaf expansion as a consequence of late spring (Fig. 2, Table 1). In the literature, most genetic studies on phenology of trees are focused on bud break [47–50] and phenology of leaf expansion has been limited to early stages [51, 52]. The fact that our putative leaf phenology controlling gene set does not overlap with these studies is probably because we compare fully expanded spring leaves (Fig. 2, Table 1, Table S1).

Summer and fall on the other hand, occur along a more compressed gene expression space, but show a considerable degree of separation along the PC2 (Fig. 1). Slight differences among the transcriptional phenotypes between the two sites could be reflecting chronically diseased states in HARV and/or latitudinal difference (Fig. 1). The unusually wet 2018 growing season appears to assign all trees to an ordination pattern that covers a wider area in the transcriptional phenotypic space (Fig. 1, shaded points). The upper and lower bounds of the ordination are set by 2018 spring and fall samples (Fig. 1).

The growing season of 2017 in HARV started with a large water deficit but followed a steady recovery until April 2018 (Fig. 3, Fig. S5). During this period trees expressed transcripts with GO terms enriched in carbohydrate: proton symporter activity (GO:005351) especially of monosaccharides (GO:0015145) including hexose transmembrane transporter activity (GO:0015145) which include Sugar Transport Protein (STP) subfamily (Fig. S8). Monosaccharide transporters represent an ancestral gene family in land plants and are adapted to overtake diverse physiological roles in vascular plants including carbohydrate partitioning [53, 54]. When overexpressed in the vasculature and mesophyll cells, STPs can increase growth and Nitrogen use in Arabidopsis [55]. In leaves, monosaccharide transporters are highly redundant but essential in non-structural carbon balance [56]. The whole-plant long distance carbon transport is mediated by sucrose. In early spring sucrose travels from roots to the buds for leaf out and is hydrolyzed to monosaccharides via invertases. Once in monosaccharide form, proton-coupled symporters commit sugar uptake [54, 56, 57]. STP-mediated transport of non-structural carbohydrates are key in survival since trees must maintain a certain reserve of mobilizable stored carbon for survival against osmotic stress and pathogen attack [58, 59]. Both stressors were present in HARV during the growing season of 2017. Plant sugar transporters are aggressive enzymes and have been shown to carry three orders of magnitude higher-affinity compared to their pathogenic counterparts and form the backbone of a defense strategy based on apoplastic sugar depletion [59]. Retrieval of carbohydrates leaked into the apoplast traveling to the phloem is a homeostatic activity during regular growth. By upregulating the monosaccharide symporter activity, trees in HARV could be showing a joint response against water deficit and pathogen attack which contrasts with SERC where these stressors didn’t exist in 2017 (Fig. S5, Fig. S7, Fig. S8).

Exceedingly wet growing season across the entire East coast of the continental US in 2018 led to simultaneous downregulation of 255 transcripts in SERC and HARV during spring-summer transition (Fig. 2, Fig. S8). These genes span 23 GO terms possibly representing decreased investment on fatty acid biosynthesis including very long chain fatty acids (VLCFAs) which are precursors for cellular membranes, wax and suberin (Fig. S8). This is intriguing since epidermal cell morphology, especially the extension of stomatal guard cell walls in the form of outer cuticular ledge, affects resistance to drought and is dependent on VLCFA deposition [60, 61]. Relaxation from expensive anabolic processes in the leaf tissue allows diversion of resources towards growth in woody tissues or greater allocation to seed storage lipids. VLCFA elongation takes place in the ER membrane through a sequential chain of four enzymes. Our analysis showed that the first enzyme in the chain, the 3-ketoacyl-CoA-synthase activity (GO:0102756) and a set of others including 3-oxo-arachidoyl-CoA synthase activity (GO:0102336) as well as 3-oxo-cerotoyl-CoA synthase activity (GO:0102337), and 3-oxo-lignoceronyl-CoA synthase activity (GO:0102338) are downregulated (Fig. S8). Chain elongation is an energy demanding anabolic process requiring 7 ATP for building the 16C palmitic acid. The reverse chemistry is also rewarding, the complete beta-oxidation of palmitic acid through Krebs Cycle, electron transport chain and glycolysis can generate 110 ATPs [62]. In less stressful times, such as in 2018, the surplus VLCFA pool could potentially be oxidized in peroxisomes to generate energy to divert growth towards non-leaf tissues. Additionally, VLCFAs are necessary for auxin transport by polar auxin influx carrier PIN1 affecting tissue patterning [63].

Finally, due to recent demographic history, even for unrelated outbred individuals, the knowledge of their population level background relatedness is important for interpreting the observed variation in gene expression [64]. New England has experienced extensive deforestation after the European settlers arrived and lost a fraction of their genetic diversity [65], [66]. Present day forests in the region are largely of second-growth that have expanded from highly bottlenecked founders or few fragmented patches that have maintained old-growth character [67]. The fraction of preserved genetic diversity appears to differ among species with different modes of pollen and seed dispersal [68–72]. The nature of the retained diversity ie. whether it is based on gene flow, emergence of novel variants through mutation and recombination as well as plasticity contribute to this picture [3, 70, 73–83]. Our kinship analysis has shown that SERC and HARV populations were genetically distinct and at the same time harbored some degree of background relatedness characteristic of second-growth forests (Fig. S4). American beech seeds are animal-dispersed. Many vertebrate consumers of its seeds include squirrels, raccoons, porcupines, foxes, black bears, bluejays, pheasants and turkeys [84–86]. Therefore, beech trees are expected to benefit from this adaptation in terms of long-distance gene flow. In fact, sporadic presence of unrelated individuals observed within the two populations could be an indicator of genetic recovery by successful long-distance seed dispersal and seedling establishment (Fig. S4).

### Conclusions

Our investigation shows that based on differential and co-expression patterns, it is possible to reconstruct the leaf physiological states of individual trees to understand forest dynamics with respect to site, year and seasonal differences. This study provides a detailed overview of transcriptomic signatures in the leaf tissue of American beech at a single-gene resolution. We show that it is possible to identify sets of differentially expressed transcripts along a time series, map them on a protein-protein interactome and extract the underpinning biology through enriched GO terms. Our genome-guided transcriptome comparison took place in a setting and at a time interval where overlapping biotic and abiotic factors perturbed the gene expression to interpretable levels and yielded explanations on the collective responses of trees to situations including recovery from drought, late spring induced phenological shift, excessive precipitation and chronic disease states. Individually, many genes may show clearly defined functions seemingly disconnected to others. However collectively, the interactions of their protein products may orchestrate a continuum of shifting phenotypes adapted to carry out life processes in optimum.

As further sequencing data from diverse tree species with more extensive time series accumulate, it will be possible to capture and understand tree responses against rare and extreme incidences together with underlying molecular dynamics shaping the phenotypic space. Here we wanted to set an example for monitoring projects such as the ForestGEO and NEON which can incorporate transcriptomics as an informative cutting-edge tool to generate temporally and spatially resolved genetic data as a legible permanent record of tree responses in forests. We hope our study serves as a template for potential scaled-up transcriptomic studies in natural forest settings.

## Materials and Methods

### Sites and species studied

The study was carried out at two ForestGEO Forest Dynamics Plots (FDPs): the 30 ha plot at Harvard Forest (HARV), Petersham, MA and the 16 ha plot at the Smithsonian Environmental Research Center (SERC), Edgewater, Maryland. HARV represents a higher latitude inland forest with relatively pristine air quality where almost all *F. grandifolia* trees are under attack from beech bark disease involving scale insects (*Cryptococcus fagisuga* and *Xylococculus betulae*) and fungal pathogen Neonectria (*N. ditissima* and *N. faginata*). HARV also experiences episodic spongy moth (*Lymantria dispar*) outbreaks. SERC represents a lower latitude, coastal forest chronically affected by daily plumes of metropolitan Washington DC area emissions. At SERC, during the time of sampling, all trees were free of beech bark disease, and no outbreak of spongy moth has been observed. The two FDPs are co-located with NEON (National Ecological Observation Network) sites, each harboring a tower equipped with instrumentation for ecological monitoring. At each FDP, 20 trees were selected according to an even size class distribution (SI Fig. S9, SI Table S1). All trees had a dendroband installed and growth measurements were taken at weekly intervals.

### Sample collection and RNA library construction

Using a modified crossbow with a fishing line and reel attachment, leaves have been felled from the canopy through spring, summer and fall between 2017 and 2018 (SI Fig. S1). For subcanopy trees we used a 12m pruner pole. Sample collections were done in the morning hours between 8:00 am to 10:30 am to avoid expression changes throughout the day. For each sample, 10 leaf discs were punched into 2ml extraction vials containing 3 steel balls of 3mm in diameter, and flash frozen in liquid nitrogen in the field (SI Fig. S2). Frozen samples were then transferred to a -80 freezer at the end of the day. RNA extractions were carried out using E.Z.N.A Plant RNA Kit by (Omega Bio-Tek, Norcross, GA, USA). Quality metrics for extracted RNA were determined using the Agilent Bioanalyzer 2100 instrument (Agilent Technologies, Santa Clara, CA, USA) and samples containing genomic DNA contaminants were treated with DNAse I (ThermoFisher Scientific, Waltham, MA, USA). Illumina TruSeq libraries were prepared for each sample RNA (Illumina Inc., San Diego, CA, USA). Libraries were indexed and pooled to be sequenced on the Illumina NovaSeq 6000 platform as 150 nucleotide long paired end reads targeting 14 million total reads per sample (Illumina Inc., San Diego, CA, USA). Sequences are uploaded to the NCBI Sequence Read Archive (SRA) with the bioproject number PRJNA630305. Sample naming convention includes Site (H for HARV, S for SERC), Year (17 for 2017, 18 for 2018), Season (1 for Spring, 2 for Summer, 3 for Fall) Species (Fg for *F. grandifolia*) and tree ID number (between 01-50).

### Assembly and annotation metrics of the draft reference American beech genome

The largest tree with dbh of 86.6 cm at the beginning of the 2017 growing season located in SERC (Sample ID SFg10 and ForestGEO FDP treetag ID 43420) served as the high molecular weight DNA source for long-read sequencing and was subsequently used to assemble the University of Connecticut (UCONN) version of the unpublished American beech draft genome. The UCONN reference assembly used long and short read sequences generated by a single Oxford Nanopore GridION run (179x coverage) at the University of California Davis (DNA Technologies and Expression Analysis Core) and Illumina HiSeq 2500 run (198x coverage) at 250 bp PE at the Harvard University (Bauer Core Facility). The UCONN reference assembly had a genome size of 443 Mbp with N50 885 Kb harboring 784 contigs including 12 putative chromosomes. Completeness assessment through Benchmarking Universal Single-Copy Orthologs (BUSCO) embryophyta set (n1614) was more than 97 % (single copy genes 88.5 %, duplicates 8.2 %, fragmented 0.7 %, missing 2.6 %) [87]. The total number of annotated genes was 48,528 with 7368 monoexonic and 41160 multiexonic genes.

### Bioinformatics

Demultiplexed reads were processed with TRIMMOMATIC for adapter removal and quality filtering of the fastq formatted raw data [88]. Trimmed reads were aligned to the *F. grandifolia* draft genome sequence using HISAT2 splice aware aligner [89]. Duplicate identification and read-ID groupings of paired-end reads were carried out with SAMBLASTER, and alignments were sorted by SAMTOOLS [90, 91]. Variants were called through FREEBAYES to generate individual VCF variant call format files for each season and consolidated into a total of 40 individuals using BCFTOOLS [92, 93]. Population genetic analysis for background genetic relatedness between FDP sites and among individual trees was done by the TASSEL5 suit through calculation of pairwise kinship matrix using centered Identity by State (IBS) genetic distances [94, 95]. The probability of IBS is based on the definition where randomly drawn alleles from two individuals at the same locus are identical. The kinship matrix was used as input to calculate Kendall’s correlation for genetic associations within and between the two FDP populations using MORPHEUS matrix visualization and analysis software (Broad Institute, Boston, MA, USA). Alignments from HISAT2 were processed by HTSEQ to generate abundance for transcripts [96]. HTSEQ counts were filtered to include top 11.5k loci with most variance. A PCA was generated from variance filtered loci using CLUSTVIS R package through ln(x) transformation with the default method of singular value decomposition and Pareto scaling [97]. We used the TUXEDO pipeline for differential gene expression analysis [98]. Based on alignments, a merged genome-guided transcriptome assembly and transcript quantification were done by STRINGTIE [99]. Per gene count data for mRNA abundance (FPKM) were calculated using the differential gene expression analysis R package BALLGOWN [100]. Differentially expressed beech transcripts passing the minimum fold change (log2fc >2.32, pval <0.01) were retained. For each year and site, differential expression analyses were carried out in the following comparative frame: Spring (condition 1) vs. Summer (condition 2) and Summer (condition 1) vs Fall (condition 2). Protein sequences of the differentially expressed transcripts were BLASTed against indexed black cottonwood (*Populus trichocarpa*) proteome with the NCBI taxon ID 3694 to serve as orthology template for downstream interactome time course analysis. Top hits were selected through VSEARCH (-ublast -evalue 1e-9 -query cov 0.9) [101]. BLAST results were used as input into the STRING database for network construction using *P. trichocarpa* interactome [102, 103]. Co-downregulated and co-expressed protein-protein interactome networks were imported into CYTOSCAPE and merged into a single network [104] (Table S1). Gene Ontology (GO) terms were compiled by using g:PROFILER by providing gene lists corresponding to *P. trichocarpa* orthologs [105]. Results generated from g:Profiler are accessible through permalinks in Table S1.

## Supporting Information

Supporting information for Sezen et al. 2023, Leaf gene expression trajectories during the growing season are consistent between sites and years in American beech. Dryad accession permalink: https://doi.org/10.5061/dryad.fxpnvx0vp.

**Table S1**

**Spreadsheet.** A multi-tab format spreadsheet containing sample list, tree size classes, genome alignments, transcript raw counts, ordination results, list of top 20 differentially expressed genes, g:Profiler output permalinks, kinship matrix, Kendall’s correlation values and an annotated list of 324 leaf phenology related gene orthologs.

**Dataset S1**

**Sequences.** A fasta file of differentially expressed 4197 beech protein sequences.

**Dataset S2**

**Variant calling.** A consolidated sorted VCF file harboring 40 trees.

**Fig. S1**

Photographic description of the crossbow sampling set up. Leaves from crowns of the tall trees were sampled using a Barnett International Youth100 draw weight 100 lbs. thrust crossbow with custom-made fishing reel attachment and variable weights on bolt tips.

**Fig. S2**

Photographic description of leaf sample collection in the field. Leaf discs have been punched out prior to freezing in liquid nitrogen.

**Fig. S3**

Percentage of samples with alignment rates of trimmed reads to draft American beech genome. Close to 95 percent of the samples had reads aligned with a rate more than 80%.

**Fig. S4**

A Kinship Analysis of SNP variation and the strength of correlated association among 40 F. grandifolia trees from HARV and SERC. (Top) The pairwise relatedness matrix is calculated based on the centered probability of Identity by State (IBS). Kinship values show a marked decrease in comparisons between the two populations. Within population comparisons show higher kinship reflecting the local founder effect. The diagonal at the bottom shows self-comparisons where kinship coefficient reaches maximum. (Bottom) Kendall’s rank correlation of association applied on kinship coefficients for within and between populations. Self-associations in the diagonal assume the value of 1.0. Within population correlation values are dominated by values above zero indicating higher genetic relatedness. Between population correlations drop to negative values indicating higher genetic differentiation. The lowest value (-0.62) was measured between HFg45 and SFg29 indicating the most distantly related trees in our sample. In HARV, three individuals HFg24, HFg25, HFg37 stand out as most distantly related within the population. Similarly, in SERC, SFg10 is the most distantly related tree within the population. The HARV tree HFg37 shows a high genetic relatedness to most trees in SERC highest value (0.74) corresponding to pairwise comparison with SFg07.

**Fig. S5**

Drought maps from 2016-2018 in the Eastern US compiled from the US Drought Monitor initiative by the National Drought Mitigation Center (NDMC) at the University of Nebraska-Lincoln, the National Oceanic and Atmospheric Administration (NOAA), and the U.S. Department of Agriculture (USDA) and the NASA Gravity Recovery and Climate Experiment (GRACE) satellite that detects changes in the Earth’s gravity field caused by water on and beneath the land surface. In 2016, New England states experienced a severe regional drought. 2017 was a recovery year from drought. The visualization switches from NDMC to GRACE in April 2017 (shown by an inset frame). In 2018 the entire US east coast experienced record breaking amounts of precipitation. The map classifications are: abnormally dry (D0), showing areas that may be going into or are coming out of drought, and four levels of drought: moderate (D1), severe (D2), extreme (D3) and exceptional (D4). The GRACE-Based Root Zone Soil Moisture Drought Indicator colors are based on a wetness percentile relative to the 1948-2012 period as baseline.

**Fig. S6**

Abundant rain across the Eastern United States in 2018 resulted in the wettest year on record. The shallow shallow groundwater on December 17, 2018, ranks among all the Decembers from 1948 to 2012. Blue areas have more abundant groundwater while orange and red areas have less compared to the mean calculated between 1948-2012 period. Washington, D.C. surpassed the previous annual rainfall record of 61.33 inches set in 1889, accumulating 64.22 inches by the date of this map. Similarly, Wilmington, North Carolina, has received 99.98 inches. Baltimore, Maryland, broke its previous record of 62.66 inches set in 2003 with 68.77 inches. The map is generated by

incorporating meteorological data with observations from NASA’s Gravity Recovery and Climate Experiment (GRACE) satellites. https://earthobservatory.nasa.gov/images/144417/soggy-2018-for-the-eastern-us

**Fig. S7**

A photographic comparison of diseased and healthy state beech trees at the two study sites. Beech bark disease has not been observed on the SERC Forest Dynamics Plot. https://forestgeo.si.edu/sites/north-america/smithsonian-environmental-research-center. The trees in HARV Forest Dynamics Plot however are on the killing front of the disease https://forestgeo.si.edu/sites/north-america/harvard-forest.

**Fig. S8**

Enriched GO terms of differentially expressed transcripts during spring-summer transitions of 2017 and 2018. Terms enriched among upregulated transcripts are shown in red. Terms related to downregulated transcripts are shown in blue.

**Fig. S9**

PCA showing tree size classes. The same PCA plot as Fig. 1 with three size classes remapped. Large (diamond), medium (cross), small (circle).

**Fig. S10**

A breakdown of differential and co-expression patterns during spring-summer, summer-fall transitions at two sites during two consecutive years mapped on the *P. trichocarpa* protein-protein interactome using the STRING database. Interactome nodes with their corresponding color combinations are shown inside the respective Venn diagrams. Due to similarity in differentially expressed genes, summer-fall transitions in two sites are represented in a single interactome panel per year. For brevity, singleton nodes are not shown.

**Fig. S11**

Temperature (C) and rainfall (mm) 10 days prior to the beginning of sampling period in HARV during the 2017 growing season denoted with Day-of-Year (DOY). Sampling days shown as horizontal bars on the insets.

**Fig. S12**

Temperature (C) and rainfall (mm) 10 days prior to the beginning of sampling period in SERC during the 2017 growing season denoted with Day-of-Year (DOY). Sampling days shown as horizontal bars on the insets.

**Fig. S13**

Temperature (C) and rainfall (mm) 10 days prior to the beginning of sampling period in HARV during the 2018 growing season denoted with Day-of-Year (DOY). Sampling days shown as horizontal bars on the insets.

**Fig. S14**

Temperature (C) and rainfall (mm) 10 days prior to the beginning of sampling period in SERC during the 2018 growing season denoted with Day-of-Year (DOY). Sampling days shown as horizontal bars on the insets.

## Competing Interests

None declared.

## Acknowledgments

We would like to thank Dr. Jill Wegrzyn for being a valuable resource person in bioinformatics and providing access to University of Connecticut’s Xanadu High-Performance Computation Cluster. We would like to extend our gratitude for Dr. Susan McEvoy and Dr. Adrian Powell for sharing the early draft beech genome sequence assemblies and for Dr. Mengmeng Lu for providing insight and discussion that has helped shape the final version of this manuscript. We are also grateful to Mark VanScoy for taking dendroband measurements in Harvard Forest. This work was funded by the US National Science Foundation (DEB-1638488).

## Author Contributions

UUS analyzed the data, prepared the figures and wrote the manuscript with input from all authors. NGS SMM SJD conceived and articulated the idea and obtained the funding. UUS JES SJW NGS collected the samples. UUS carried out the molecular lab work. NGS coordinated the sequencing, data curation, analytical support. SJW SJD SMM JES NGS contributed to the conceptualization, writing, review and proofreading, JES SJD NGS helped with the project coordination and implementation.

## Notes

https://doi.org/10.5061/dryad.fxpnvx0vp

### Competing Interest Statement

The authors have declared no competing interest.

### Summary of Updates

I simplified Fig. 2 by removing the interactome section into the supplementary (S10) but retained the Venn diagrams. I reduced the scale of Fig. 3 and included dates of first leaf flush for Fagus. I added new figures to the supplementary (S11-14) that show temperature and precipitation data at least 10 days prior to the beginning of each sampling overlaid on the precipitation data for each growing season. I also added tree dbh in the supplementary table and mapped size classes onto the PCA as a supplementary figure (S9). I amended a new subsection to the Methods providing some basic metrics for the draft reference genome assembly used in this study. With the inclusion of suggested references the total number of citations is now 105.

