## Supplementary Figures for "Leaf gene expression trajectories during the growing season are consistent between sites and years in American beech"

**This PDF file includes:**

Supplementary text
Figs. S1 to S14

### Supporting Information Text

**Leaf gene expression trajectories during the growing season are consistent between sites and years in American beech.**

**U.Uzay Sezen, Jessica Shue, Samantha J. Worthy, Stuart J. Davies, Sean McMahon, Nathan G. Swenson.**

**Corresponding Author: Uğur Uzay Sezen**
**.**

**This PDF file includes:**
**Supplementary text**
**Figs. S1 to S8.**

**SI Figure S1.** Photographic description of the crossbow-mediated sampling set up. Leaves from crowns of the tall trees were sampled using a Barnett International Youth100 draw weight 100 lbs. thrust crossbow with custom-made fishing reel attachment and variable weights on bolt tips.

**NCBI-SRA accessions of the American beech transcriptome set.** NCBI-SRA accession number PRJNA630305.

**Permalink for Dryad accession.** : https://doi.org/10.5061/dryad.fxpnvx0vp.

**Spreadsheet.** A multi-tab format spreadsheet containing sample list, tree size classes, genome alignments, transcript raw counts, ordination results, list of top 20 differentially expressed genes, g:Profiler output permalinks and kinship matrix. **Sequences.** A fasta file of differentially expressed 4197 beech protein sequences. **Variant calling.** A consolidated sorted VCF file harboring 40 trees.

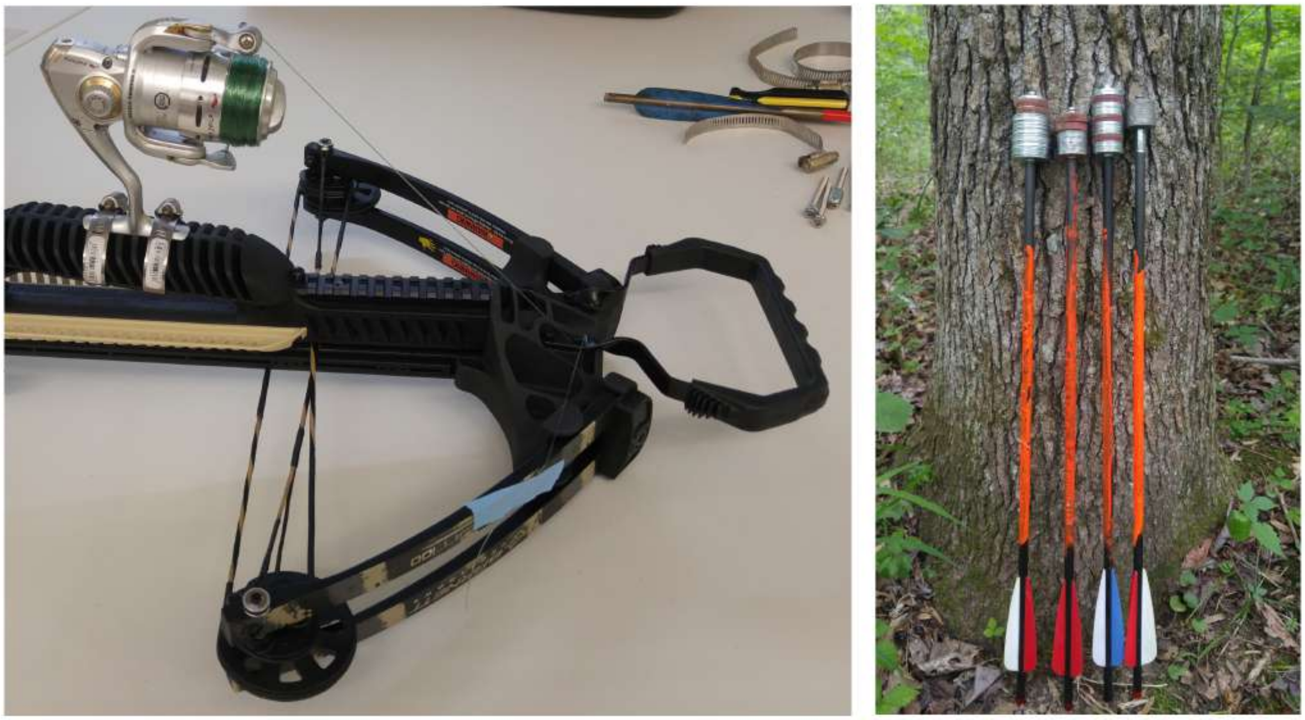

**Fig. S1.** Photographic description of the crossbow-mediated sampling set up. Leaves from crowns of the tall trees were sampled using a Barnett International Youth100 draw weight 100 lbs. thrust crossbow with custom-made fishing line and reel attachment and variable weights on bolt tips.

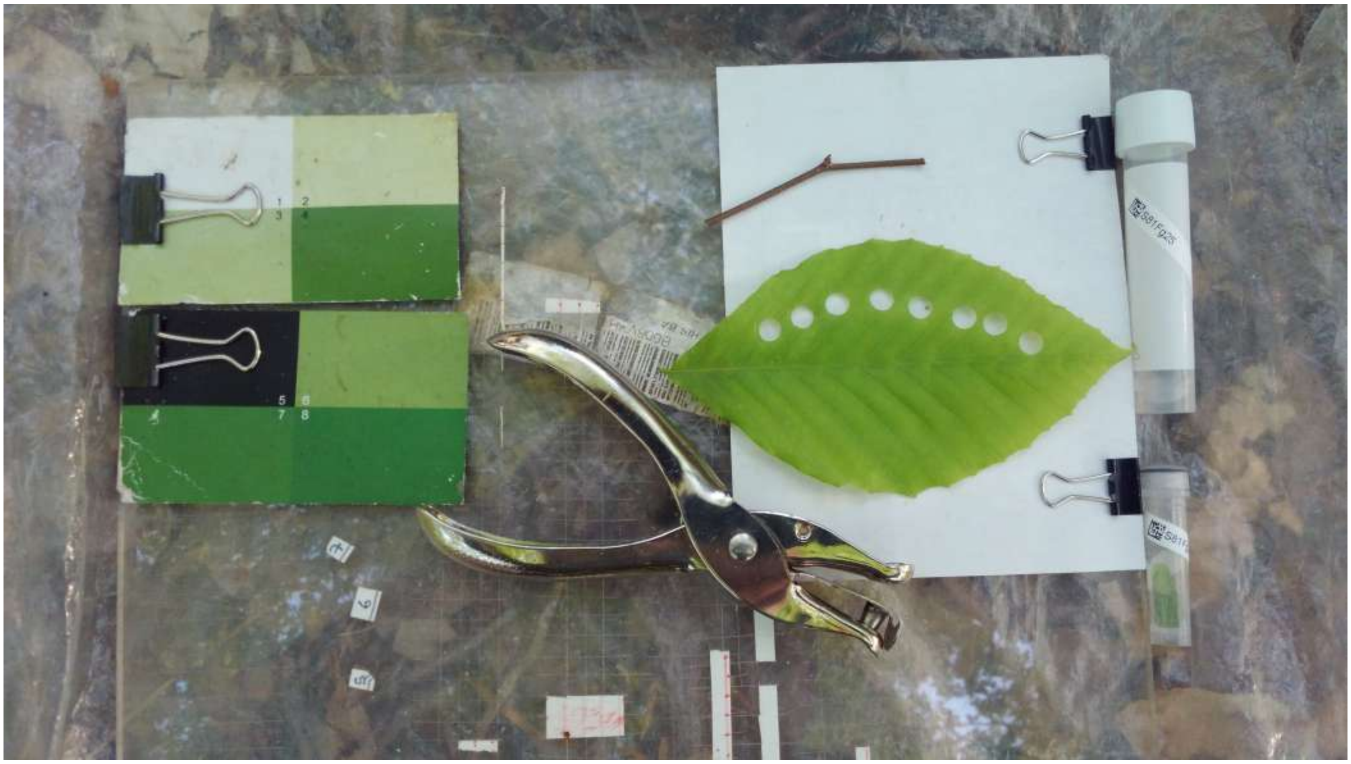

**Fig. S2.** Photographic description of leaf sample collection in the field. Leaf discs have been punched out prior to freezing in liquid nitrogen.

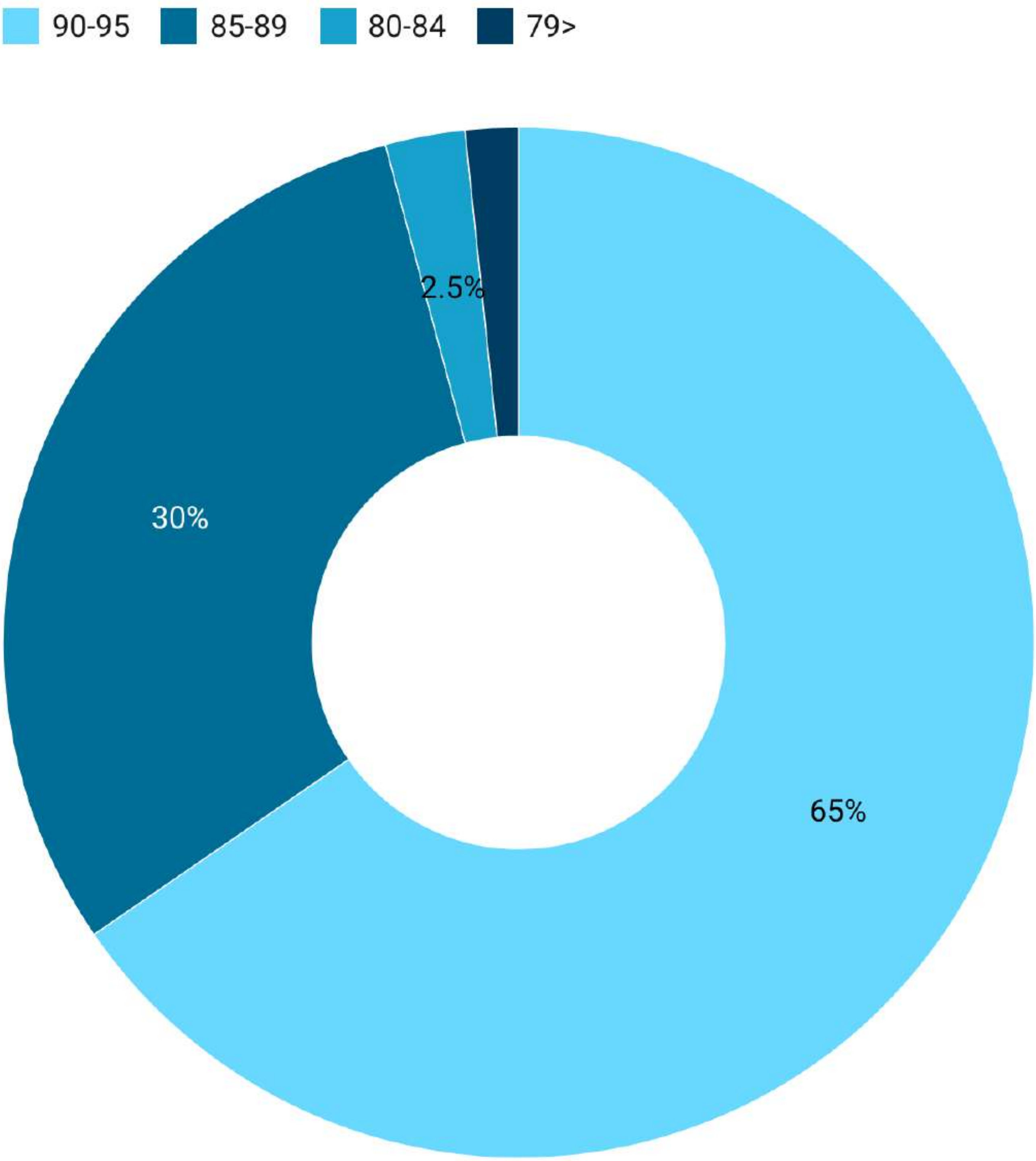


**Fig. S3.** Percentage of samples with alignment rates of trimmed reads to draft American beech genome. Close to 95 percent of the samples had reads aligned with a rate more than 80%. .

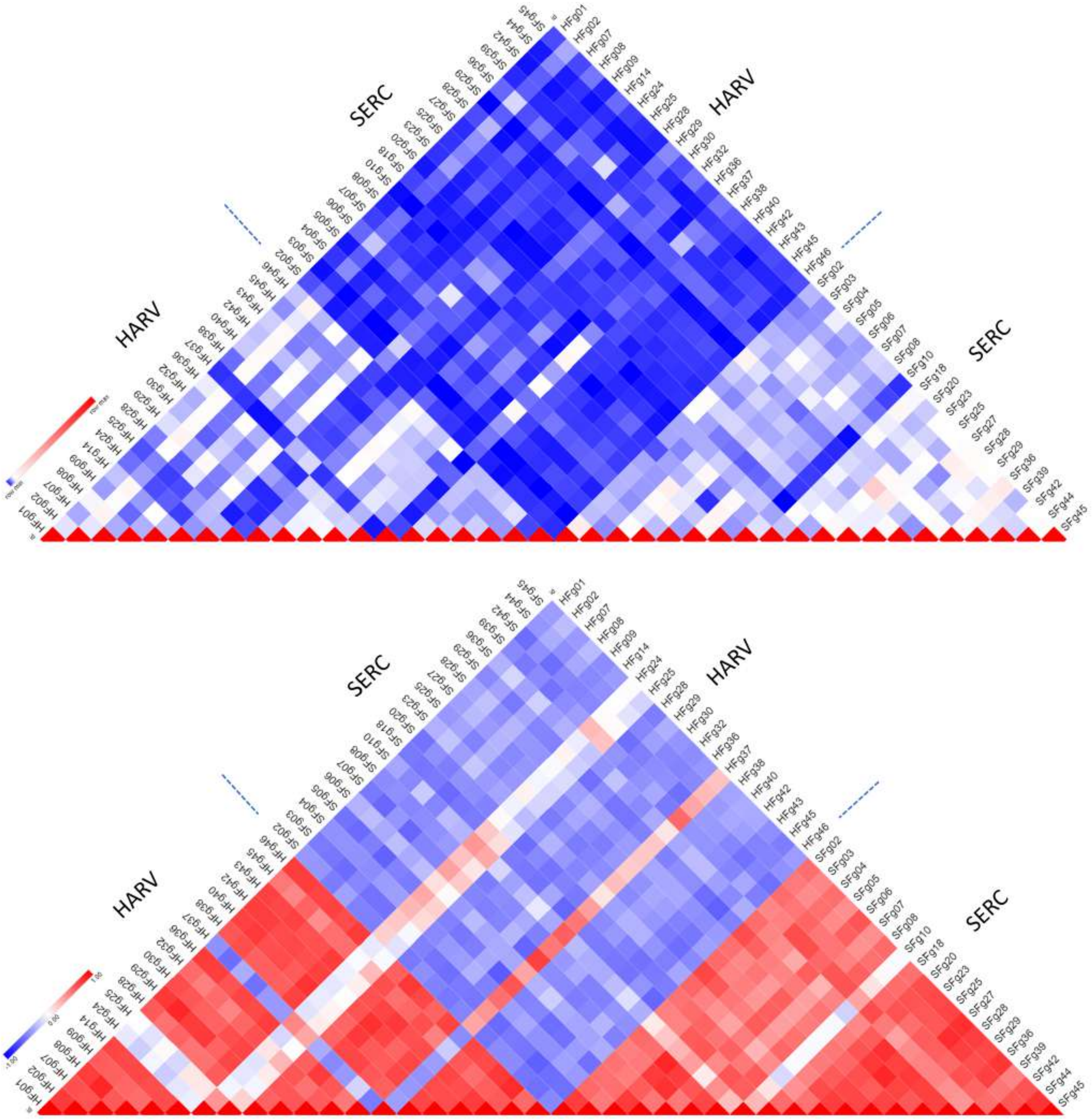


**Fig. S4. A kinship analysis of SNP variation and the strength of correlated association among 40 *F. grandifolia* trees from HARV and SERC.** (Top) The pairwise relatedness matrix is calculated based on the centered probability of Identity by State (IBS). Kinship values show a marked decrease in comparisons between the two populations. Within population comparisons show higher kinship reflecting the local founder effect. The diagonal at the bottom shows self-comparisons where kinship coefficient reaches maximum. (Bottom) Kendall's rank correlation of association applied on kinship coefficients for within and between populations. Self-associations in the diagonal assume the value of 1.0. Within population correlation values are dominated by values above zero indicating higher genetic relatedness. Between population correlations drop to negative values indicating higher genetic differentiation. The lowest value (-0.62) was measured between HFg45 and SFg29 indicating the most distantly related trees in our sample. In HARV, three individuals HFg24, HFg25, HFg37 stand out as most distantly related within the population. Similarly, in SERC, SFg10 is the most distantly related tree within the population. The HARV tree HFg37 shows a high genetic relatedness to most trees in SERC highest value (0.74) corresponding to pairwise comparison with SFg07.

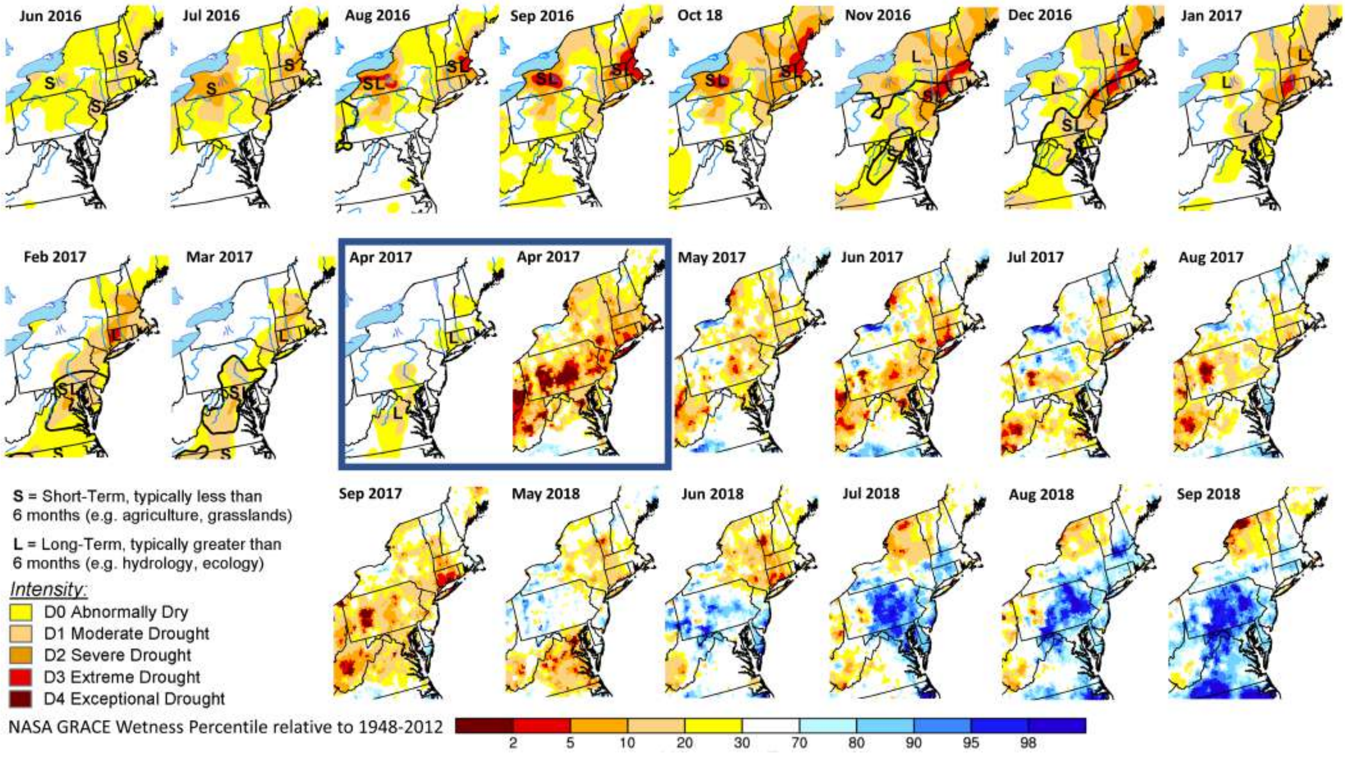


**Fig. S5.** Drought maps from 2016-2018 in the Eastern US compiled from the US Drought Monitor initiative by the National Drought Mitigation Center (NDMC) at the University of Nebraska-Lincoln, the National Oceanic and Atmospheric Administration (NOAA), and the U.S. Department of Agriculture (USDA) and the NASA Gravity Recovery and Climate Experiment (GRACE) satellite that detects changes in the Earth's gravity field caused by water on and beneath the land surface. In 2016, New England states experienced a severe regional drought. 2017 was a recovery year from drought. The visualization switches from NDMC to GRACE in April 2017 (shown by an inset frame). In 2018 the entire US east coast experienced record breaking amounts of precipitation. The map classifications are: abnormally dry (D0), showing areas that may be going into or are coming out of drought, and four levels of drought: moderate (D1), severe (D2), extreme (D3) and exceptional (D4). The GRACE-Based Root Zone Soil Moisture Drought Indicator colors are based on a wetness percentile relative to the 1948-2012 period as baseline.

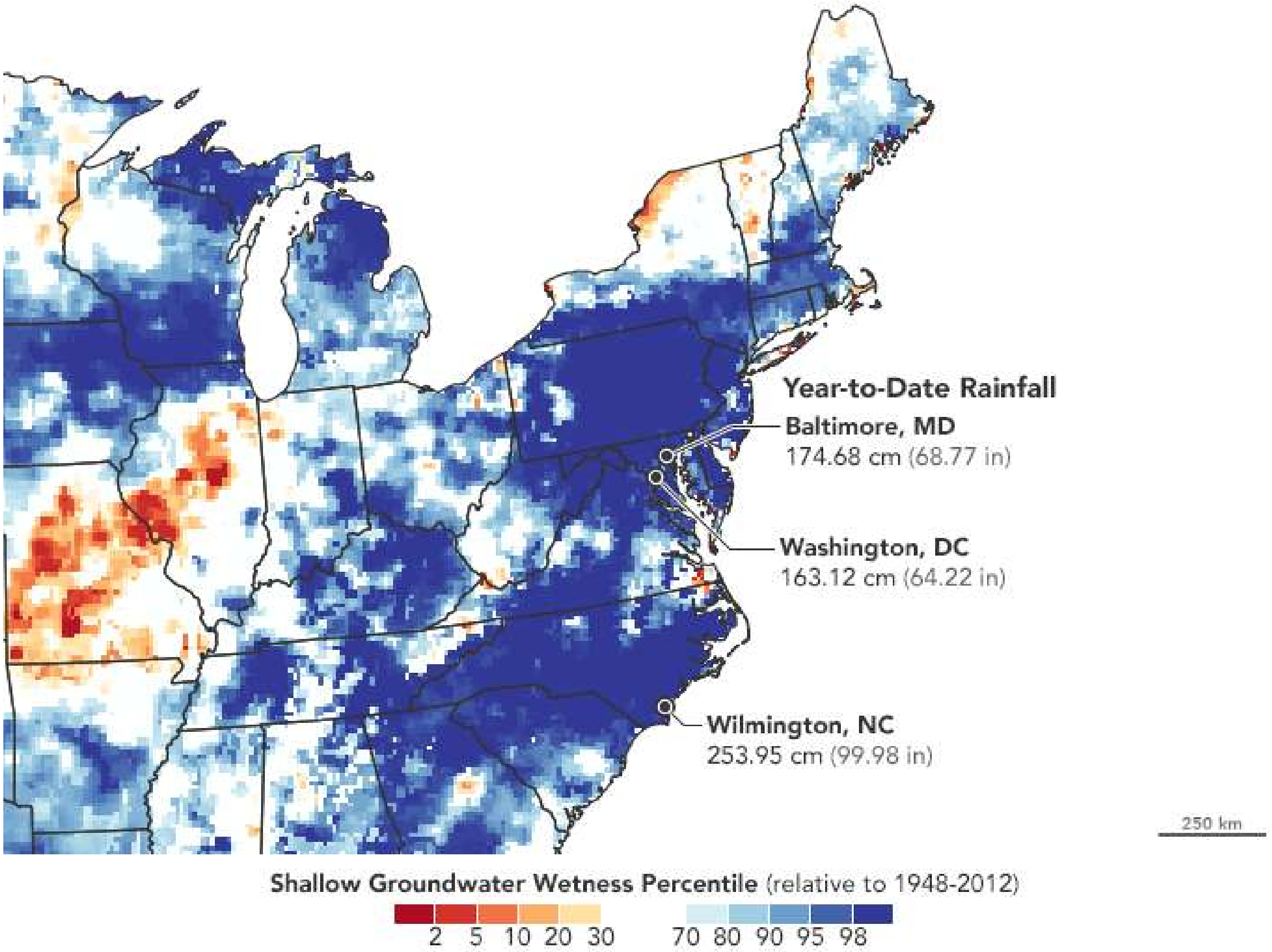


**Fig. S6.** Abundant rain across the Eastern United States in 2018 resulted in the wettest year on record. The shallow shallow groundwater on December 17, 2018, ranks among all the Decembers from 1948 to 2012. Blue areas have more abundant groundwater while orange and red areas have less compared to the mean calculated between 1948-2012 period. Washington, D.C. surpassed the previous annual rainfall record of 61.33 inches set in 1889, accumulating 64.22 inches by the date of this map. Similarly, Wilmington, North Carolina, has received 99.98 inches. Baltimore, Maryland, broke its previous record of 62.66 inches set in 2003 with 68.77 inches. The map is generated by incorporating meteorological data with observations from NASA's Gravity Recovery and Climate Experiment (GRACE) satellites. https://earthobservatory.nasa.gov/images/144417/soggy-2018-for-the-eastern-us

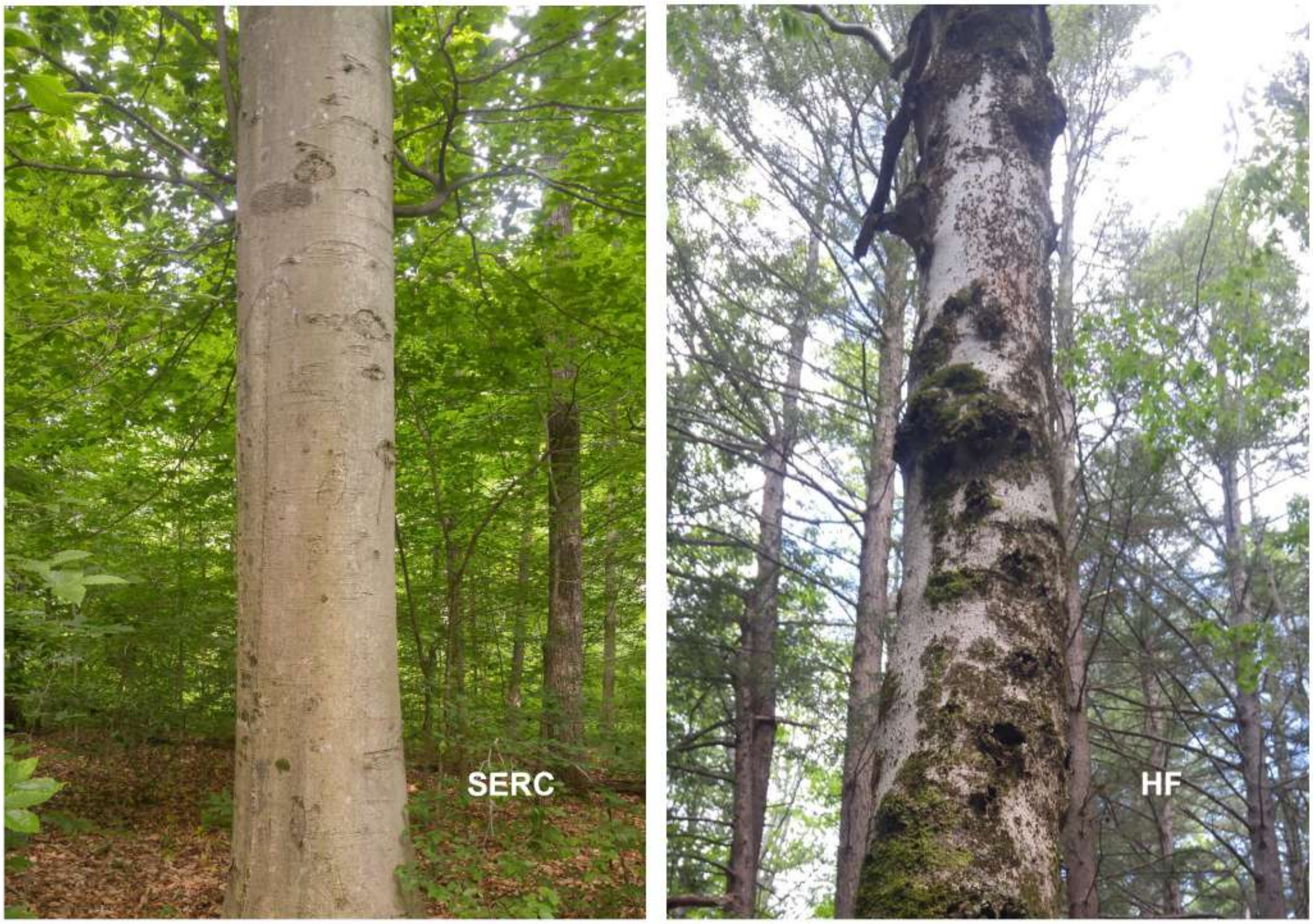


**Fig. S7.** A photographic comparison of diseased and healthy state beech trees at the two study sites. Beech bark disease has not been observed on the SERC Forest Dynamics Plot. https://forestgeo.si.edu/sites/north-america/smithsonian-environmental-research-center. The trees in HARV Forest Dynamics Plot however are on the killing front of the disease https://forestgeo.si.edu/sites/north-america/harvard-forest.

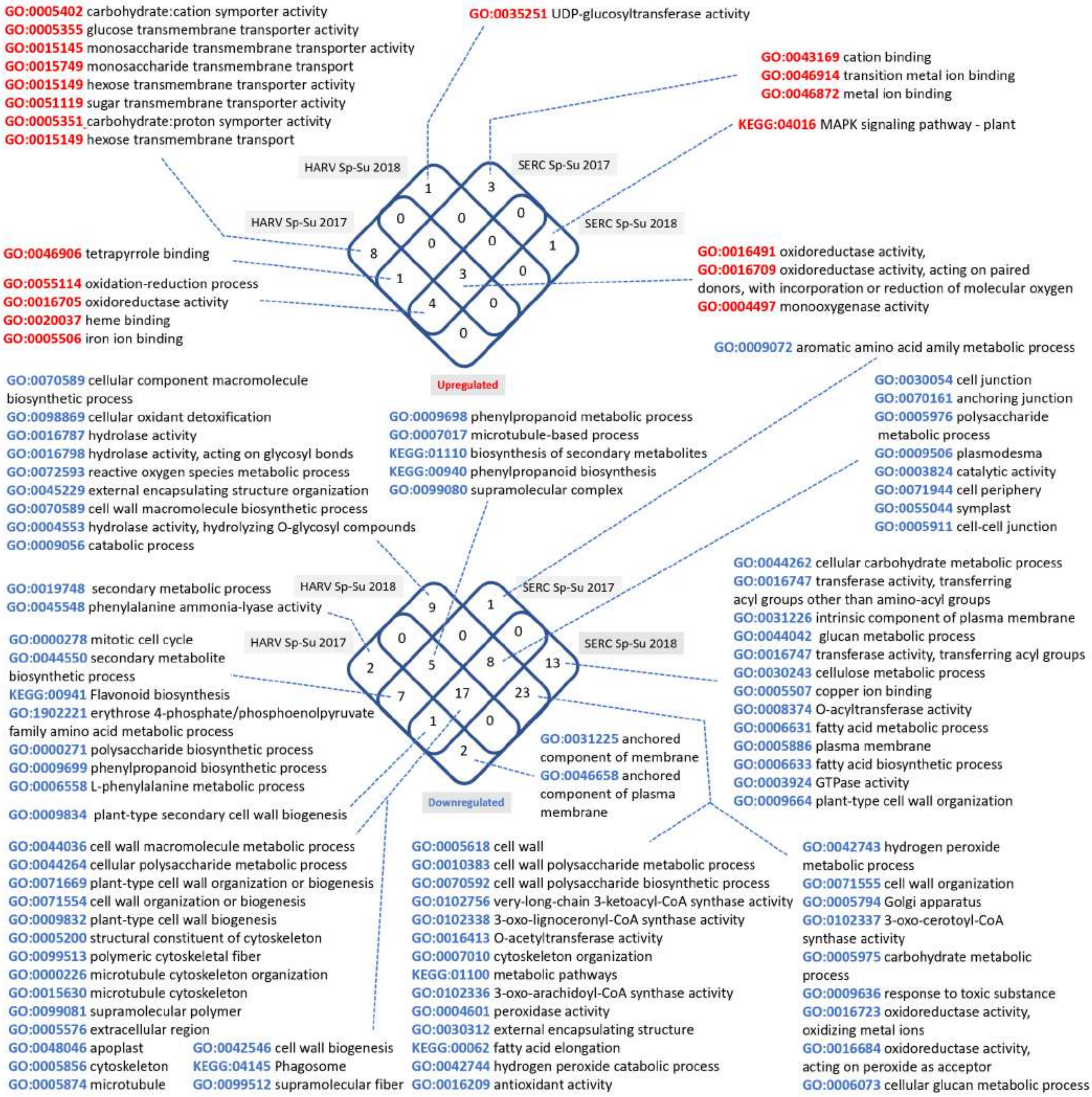


**Fig. S8. Enriched GO terms of differentially expressed transcripts during spring-summer transitions of 2017 and 2018.** Terms enriched among upregulated transcripts are shown in red. Terms related to downregulated transcripts are shown in blue.

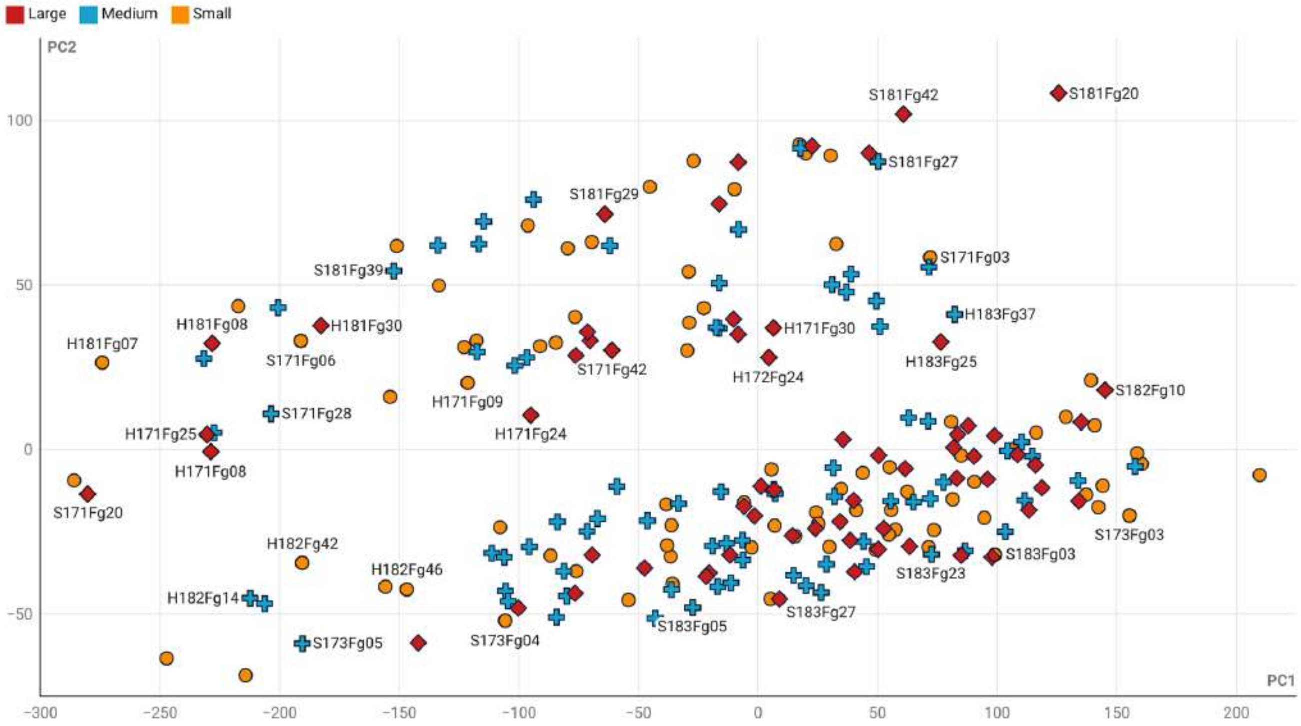


**Fig. S9. PCA showing tree size classes.** The same PCA plot as Fig. 1 with three size classes remapped. Large (diamond), medium (cross), small (circle)

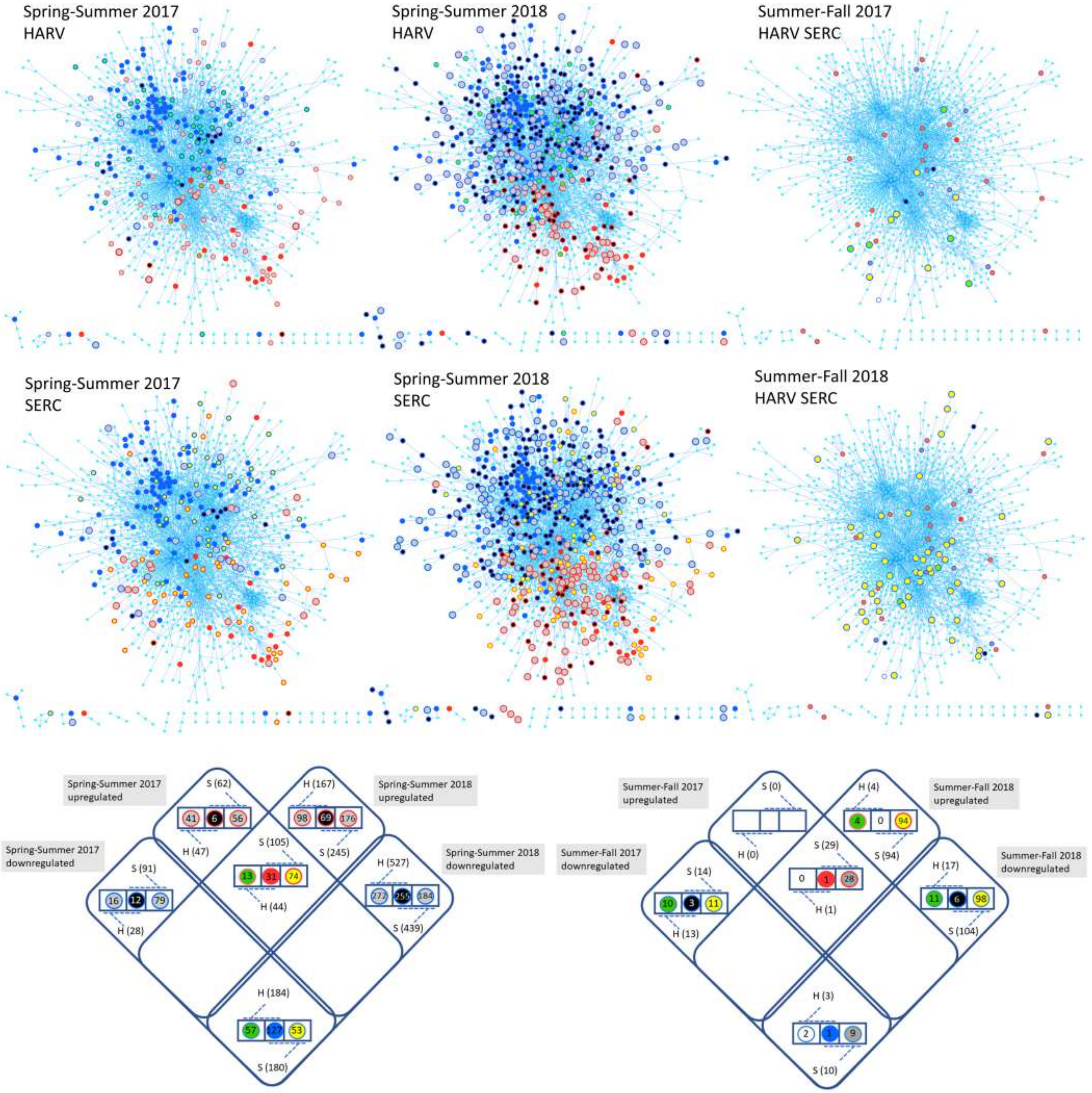


**Fig. S10. A breakdown of differential and co-expression patterns during spring-summer, summer-fall transitions at two sites during two consecutive years mapped on the *P. trichocarpa* protein-protein interactome using the STRING database.** Interactome nodes with their corresponding color combinations are shown inside the respective Venn diagrams. Due to similarity in differentially expressed genes, summer-fall transitions in two sites are represented in a single interactome panel per year. For brevity, singleton nodes are not shown.

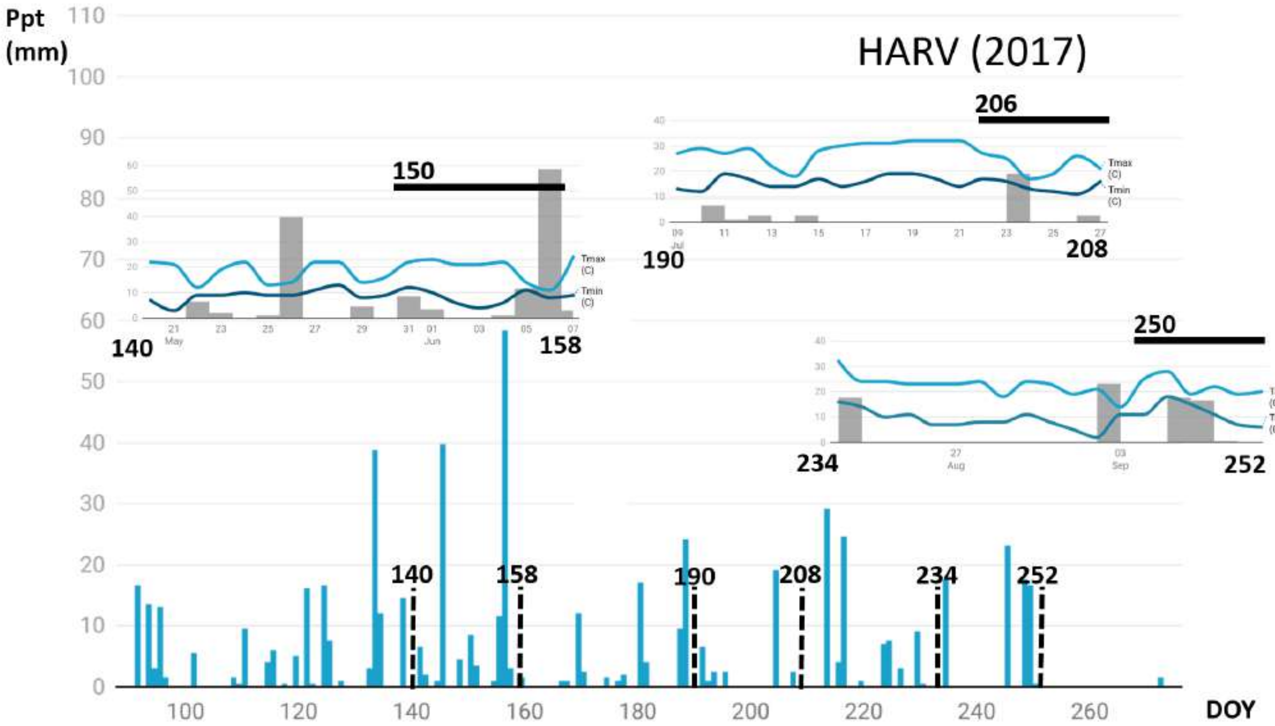


**Fig. S11. Temperature (C) and rainfall (mm) 10 days prior to the beginning of sampling period in HARV during the 2017 growing season denoted with Day-of-Year (DOY).** Sampling days shown as horizontal bars on the insets.

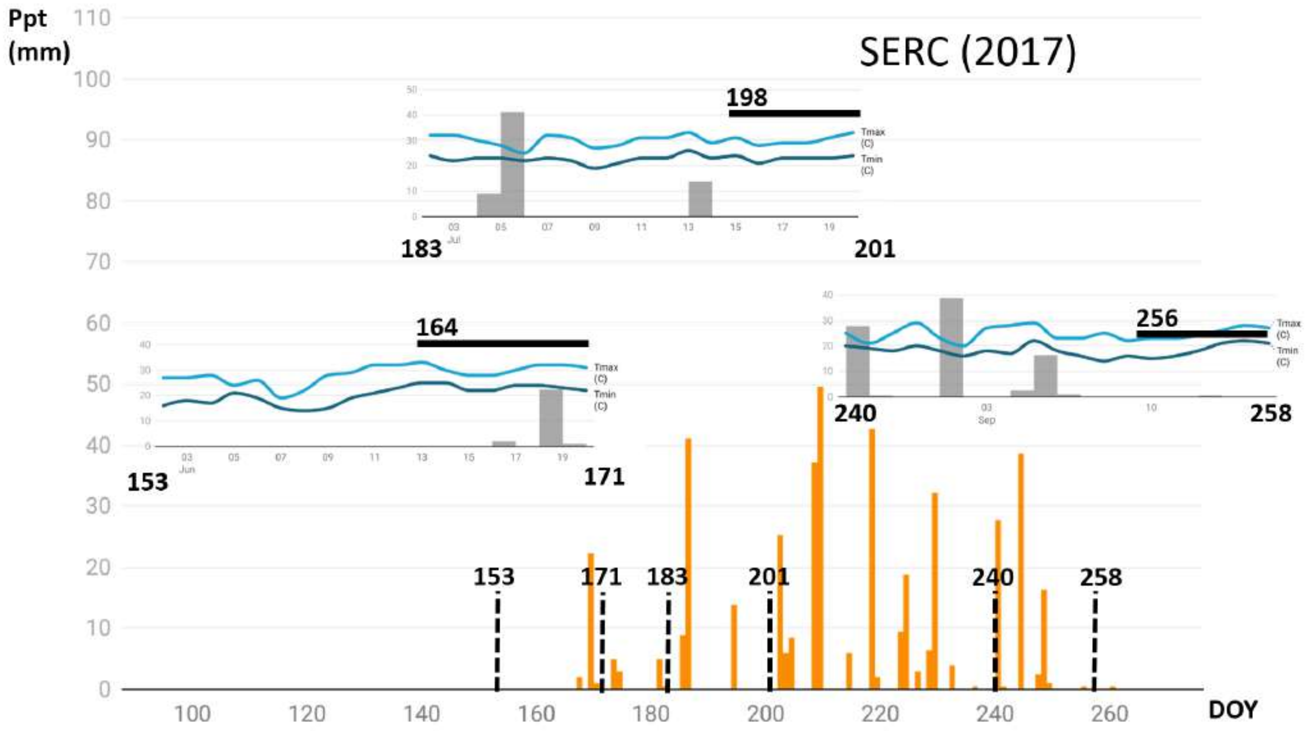


**Fig. S12. Temperature (C) and rainfall (mm) 10 days prior to the beginning of sampling period in SERC during the 2017 growing season denoted with Day-of-Year (DOY).** Sampling days shown as horizontal bars on the insets.

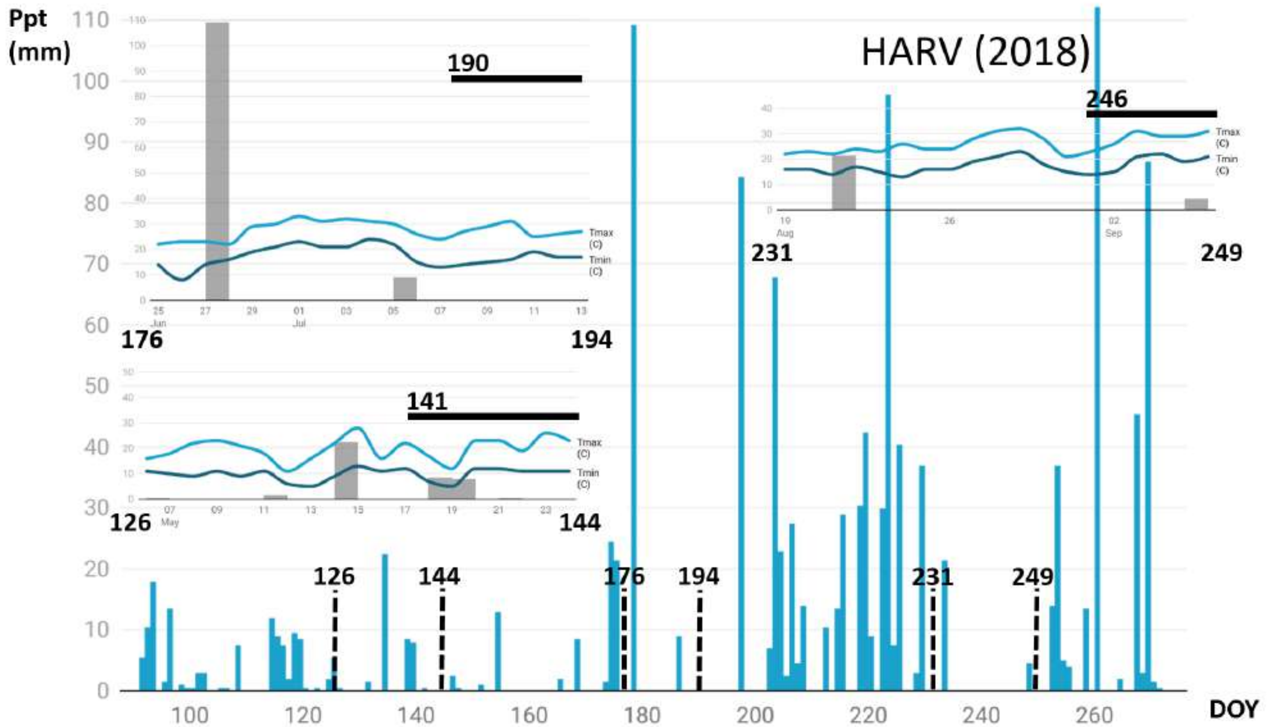


**Fig. S13. Temperature (C) and rainfall (mm) 10 days prior to the beginning of sampling period in HARV during the 2018 growing season denoted with Day-of-Year (DOY).** Sampling days shown as horizontal bars on the insets.

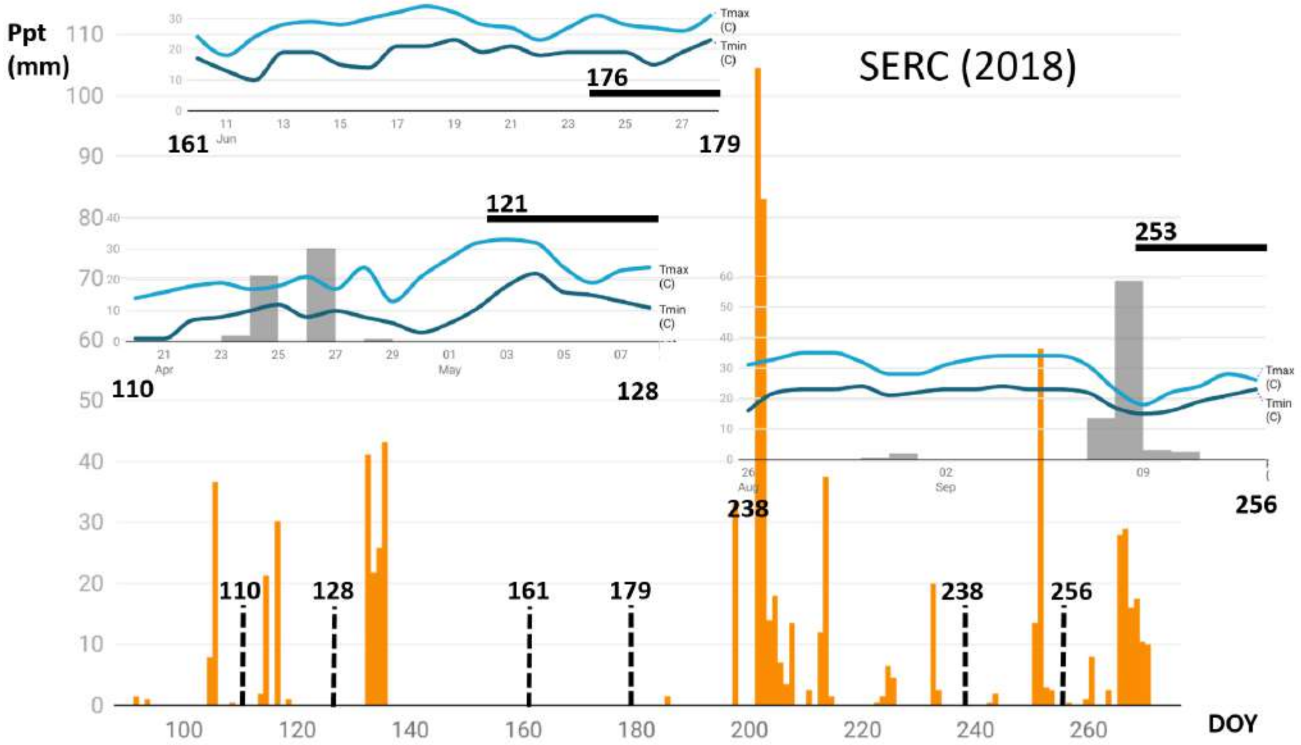


**Fig. S14. Temperature (C) and rainfall (mm) 10 days prior to the beginning of sampling period in SERC during the 2018 growing season denoted with Day-of-Year (DOY).** Sampling days shown as horizontal bars on the insets.
